## Supplementary material for "SOX7: Autism Associated Gene Identified by Analysis of Multi-Omics Data": No link

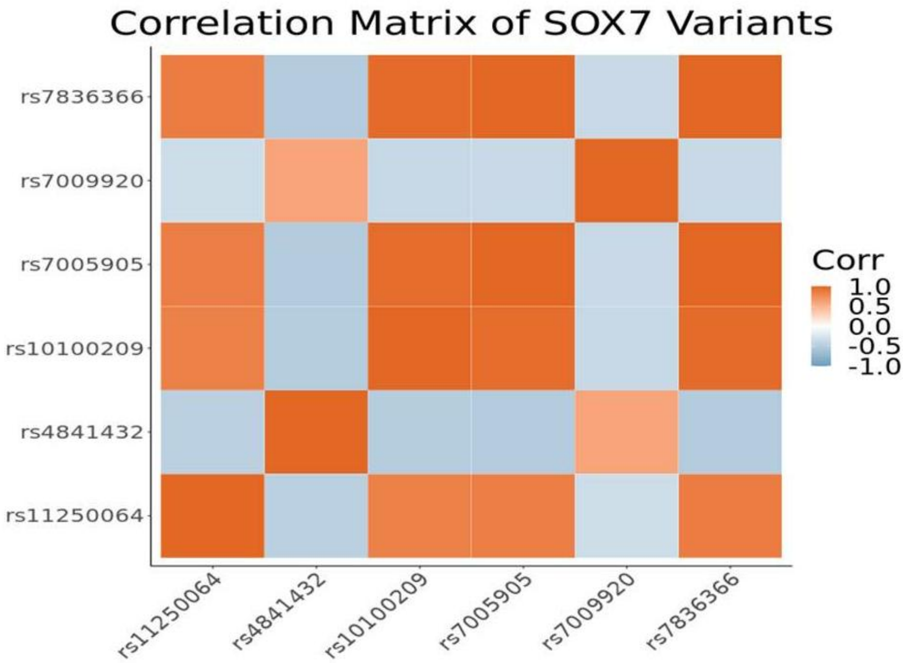


**Supplementary Figure 1. Heatmap of the correlation between variants in SOX7.** rs7005905 and rs7836366, rs10100209 and rs7836366, and rs10100209 and rs7005905 have strong positive linkage disequilibrium (LD) ($\rho$>0.5); rs4841432 has negative LD with other variants except for rs7009920.

**Supplementary Table 1. Characteristics of participants and gene SOX7 expression status in GSE211154 RNA-seq data.**

| Variable | Autism Cases (n=20) | Controls (n=19) | p-value* |
|---|---|---|---|
| **Age at dealth (Year) (Median [Inter-quartile range(IQR)])** | 17.5 (11.5) | 20 (11) | 0.44 |
| **Postmortem interval (IQR hour)** | 22.5 (5.8) | 15 (8.5) | 0.06 |
| **Sex (Male No. [%])** | 17 (85%) | 16 (84%) | 0.95 |
| **Race (No. [%])** | | | |
| White | 16 (80%) | 12 (63.2%) | 0.25 |
| Black | 4 (20%) | 7 (36.8%) | |
| **SOX7 in all samples (Median [IQR])** | 22.5 (14.25) | 16 (11) | |
| Low expression (counts ≤median) (No. [%]) | 8 (40%) | 13 (68.4%) | 0.08 |
| High expression (count>median) (No. [%]) | 12 (60%) | 6 (31.6%) | |
| **SOX7 in white samples (Median [IQR])** | 21 (18.5) | 14 (4.3) | |
| Low expression (counts ≤median) (No. [%]) | 6 (37.5%) | 10 (83.3%) | 0.03 |
| High expression (count>median) (No. [%]) | 10 (62.5%) | 2 (16.7%) | |

Note: *p-value of Z test for each predictor is obtained from univariate logistic regression.

| **Supplementary Table 2: 2-SMR Results between ASD (Outcome) and *SOX7* Expression (Exposure)** | | | |
|---|---|---|---|
| **SOX7 Expressing Tissue** | **Beta** | **SE** | **P-value[1]** |
| Adipose Subcutaneous | 0.0387 | 0.0649 | 5.51e-01 |
| Adipose Visceral Omentum | 0.2084 | 0.1119 | 6.27e-02 |
| Adrenal Gland | 0.2808 | 0.0567 | **7.31e-07** |
| Artery Coronary | -0.3076 | 0.1021 | 2.59e-03 |
| Artery Tibial | 0.2372 | 0.1375 | 8.46e-02 |
| Brain Amygdala | 0.0275 | 0.0514 | 5.93e-01 |
| Brain Anterior cingulate cortex BA24 | -0.0183 | 0.0614 | 7.65e-01 |
| Brain Caudate basal ganglia | -0.0101 | 0.0708 | 8.86e-01 |
| Brain Cerebellar Hemisphere | 0.1054 | 0.0304 | **5.31e-04** |
| Brain Frontal Cortex BA9 | 0.0097 | 0.0381 | 7.98e-01 |
| Brain Hypothalamus | 0.1799 | 0.0514 | **4.71e-04** |
| Brain Putamen basal ganglia | -0.0225 | 0.0753 | 7.65e-01 |
| Brain Spinal cord cervical c-1 | 0.1997 | 0.0601 | **8.84e-04** |
| Brain Substantia nigra | -0.1037 | 0.0380 | 6.39e-03 |
| Cells Cultured fibroblasts | 0.1299 | 0.0582 | 2.58e-02 |
| Colon Sigmoid | 0.0453 | 0.0936 | 6.29e-01 |

**Supplementary Table 2: 2-SMR Results between ASD (Outcome) and *SOX7* Expression (Exposure)**

| SOX7 Expressing Tissue | Beta | SE | P-value[1] |
|---|---|---|---|
| Esophagus Gastroesophageal Junction | 0.1054 | 0.0837 | 2.08e-01 |
| Esophagus Muscularis | 0.0192 | 0.1066 | 8.57e-01 |
| Lung | -0.5016 | 0.1290 | **1.01e-04** |
| Muscle Skeletal | 0.3941 | 0.1186 | **8.89e-04** |
| Nerve Tibial | -0.1195 | 0.0724 | 9.89e-02 |
| Ovary | -0.1701 | 0.0731 | 1.99e-02 |
| Pancreas | -0.0854 | 0.0467 | 6.74e-02 |
| Prostate | -0.1206 | 0.0573 | 3.53e-02 |
| Skin Not Sun Exposed Suprapubic | 0.2315 | 0.1012 | 2.21e-02 |
| Skin Sun Exposed Lower leg | 0.1934 | 0.0845 | 2.21e-02 |
| Small Intestine Terminal Ileum | -0.1725 | 0.1531 | 2.60e-01 |
| Stomach | 0.1453 | 0.1155 | 2.08e-01 |
| Testis | 0.1256 | 0.0482 | 9.10e-03 |
| Thyroid | 0.0406 | 0.0713 | 5.69e-01 |

[1]Bold values represent Bonferroni significant p-values ($p \leq 0.0017$)

| **Supplementary Table 3: 2-SMR Results between ASD (Outcome) and *SOX7* Expression (Exposure) (EUR GTEx Samples Only)** | | | |
|---|---|---|---|
| **SOX7 Expressing Tissue** | **Beta** | **SE** | **P-value[1]** |
| Adipose Visceral Omentum | 0.2158 | 0.0994 | 2.99e-02 |
| Adrenal Gland | -0.1299 | 0.0525 | 1.34e-02 |
| Artery Aorta | 0.0807 | 0.0945 | 3.93e-01 |
| Artery Tibial | 0.1992 | 0.1155 | 8.46e-02 |
| Brain Amygdala | -0.1670 | 0.0423 | **7.74e-05** |
| Brain Anterior cingulate cortex BA24 | -0.0463 | 0.0576 | 4.21e-01 |
| Brain Caudate basal ganglia | -0.0104 | 0.0729 | 8.86e-01 |
| Brain Cerebellar Hemisphere | 0.0989 | 0.0285 | **5.31e-04** |
| Brain Frontal Cortex BA9 | 0.0102 | 0.0399 | 7.98e-01 |
| Brain Putamen basal ganglia | 0.0057 | 0.0545 | 9.16e-01 |
| Brain Spinal cord cervical c-1 | -0.1655 | 0.0645 | 1.03e-02 |
| Brain Substantia nigra | -0.0904 | 0.0318 | 4.46e-03 |
| Colon Sigmoid | 0.0458 | 0.0946 | 6.29e-01 |
| Colon Transverse | -0.2608 | 0.0981 | 7.83e-03 |
| Esophagus Gastroesophageal Junction | 0.2045 | 0.0784 | 9.10e-03 |

| **Supplementary Table 3: 2-SMR Results between ASD (Outcome) and *SOX7* Expression (Exposure) (EUR GTEx Samples Only)** | | | |
|---|---|---|---|
| **SOX7 Expressing Tissue** | **Beta** | **SE** | **P-value[1]** |
| Esophagus Muscularis | 0.0230 | 0.0889 | 7.96e-01 |
| Heart Atrial Appendage | -0.1378 | 0.0814 | 9.07e-02 |
| Liver | -0.0677 | 0.0316 | 3.21e-02 |
| Muscle Skeletal | -0.0887 | 0.1119 | 4.28e-01 |
| Nerve Tibial | 0.1512 | 0.0780 | 5.26e-02 |
| Pancreas | 0.0732 | 0.0394 | 6.34e-02 |
| Skin Not Sun Exposed Suprapubic | 0.2111 | 0.0977 | 3.07e-02 |
| Skin Sun Exposed Lower leg | 0.0503 | 0.0936 | 5.91e-01 |
| Small Intestine Terminal Ileum | -0.0639 | 0.0836 | 4.45e-01 |
| Spleen | 0.0120 | 0.0552 | 8.28e-01 |
| Stomach | -0.0746 | 0.1116 | 5.04e-01 |
| Testis | 0.1154 | 0.0450 | 1.04e-02 |
| Thyroid | 0.0122 | 0.0660 | 8.53e-01 |

[1]Bold values represent Bonferroni significant p-values ($p \leq 0.0018$)
